# Fast Optoacoustic Mesoscopy of Microvascular Endothelial Dysfunction in Cardiovascular Risk and Disease

**DOI:** 10.1101/2021.09.28.462150

**Authors:** Hailong He, Angelos Karlas, Nikolina-Alexia Fasoula, Michael Kallmayer, Juan Aguirre, Hans-Henning Eckstein, Vasilis Ntziachristos

## Abstract

Microvascular endothelial dysfunction (ED) precedes the ED in larger arteries and is an early marker of cardiovascular disease (CVD). While precise assessment of microvascular ED could thus be used for the early detection and risk stratification of CVD, detailed interrogation of skin microvascular ED is limited by the technology available. Herein, we applied a novel approach for the non-invasive assessment of skin microvascular ED by developing fast plane raster-scan optoacoustic mesoscopy (FP-RSOM) to visualize and quantify skin microvasculature perfusion changes during post-occlusive hyperemia (PORH) tests. We combined static three-dimensional RSOM imaging with fast dynamic FP-RSOM measurements (1 frame / second) in human skin *in vivo*, which allowed for the first time to fully visualize the cutaneous microvascular response and further quantify changes of individual vessel diameter, total blood volume and vessel density during the PORH process. We further computed biomarkers from FP-RSOM images to quantify skin endothelial function within different skin layers as a function of skin depth, while conventional approaches mainly measure overall changes within sampled tissue volumes. FP-RSOM applied on smokers and patients with CVD showed clear ED in both groups compared to healthy volunteers. Moreover, FP-RSOM imaging showed higher sensitivity in quantifying the effects of smoking and CVD on skin microvascular endothelial function compared to clinically used laser Doppler flowmetry and tissue spectrometry (O2C). Our study introduces FP-RSOM as a novel tool to visualize and quantify skin microvascular ED as an early marker for the diagnostics and monitoring of cardiovascular risk and disease.

## Introduction

Endothelial dysfunction (ED) is a systemic condition that affects both the macro- and microvasculature and precedes cardiovascular disease (CVD), including coronary artery disease (CAD) and heart failure (HF). Apart from overt CVD, ED is also associated with major cardiovascular risk factors (CVRF), such as age, smoking, dyslipidaemia, hypertension, and diabetes ^1-5^. ED is commonly assessed non-invasively by means of flow-mediated dilation (FMD) or the post-occlusive reactive hyperemia (PORH) test where a large peripheral artery of the arm (e.g., brachial, radial) is measured before, during, and following a transient period of cuff-induced arterial occlusion. The FMD percentage of the measured arterial diameter in response to abrupt cuff-deflation or increase in blood flow-induced wall shear stress reflects the functional level of the endothelium and is usually assessed using ultrasound imaging. While the macrovascular endothelium is frequently assessed in clinical practice and relevant studies, recent evidence suggests that the study of microvascular ED is of equal or even greater importance ^6,7^.

In particular, microvascular (diameter < 150-200 um) ED is not only a marker of CVD and its progression ^6,7^ but it also precedes the development of macrovascular disease and associated CVRF ^8-11^. Skin microvascular ED is implicated in CAD and HF and manifests in subjects with CVRF, such as hypertension, dyslipidemia and smoking ^12^. In analogy to the macrovascular ED testing described above, skin microvascular ED is usually assessed by the corresponding cuff-induced response of the microvasculature within the skin by means of techniques which monitor the local skin perfusion over time, such as laser Doppler flowmetry (LDF) or near-infrared spectroscopy (NIRS). Thus, skin microvasculature may serve as an accessible vascular bed for the convenient non-invasive investigation of systemic ED aiming at the early detection and subsequent CVD progression and therapy monitoring ^7,11^. Nevertheless, despite its established importance, microvascular ED is not routinely studied in everyday practice partly due to drawbacks of existing measurement or imaging techniques applied in ED assessment ^3,13-16^.

Peripheral arterial tonometry (PAT) has been employed to quantify endothelial function by measuring the pulse volume amplitude (PVA) in the fingertip ^17,18^. However, PAT readouts consist of 1D-signals representing rough tissue volume changes, which may indirectly reflect but do not directly resolve information about the changes occurring within different skin layers at high resolution ^3^. The PAT signal can be easily influenced by variable non-endothelial factors ^3^. Near-infrared spectroscopy (NIRS) has been extensively used to monitor muscle perfusion and oxygenation during endothelial function tests. However, the tissue penetration depth of NIRS of up to 1 cm makes it highly challenging to separate signals from the skin and those from deeper tissues (e.g., subcutaneous fat, muscle), while its high sensitivity to light scattering limits the achieved resolution to ≈ 5-10 mm and the accuracy of NIRS can be affected in the estimation of optical path length for light transmitted through tissue ^16^. Similarly, single-point LDF provides an estimate of skin perfusion within small tissue volumes of ≈ 1 mm^3^ at depths of 1–1.5 mm into the dermis by means of abstract 1D signal recordings ^19,20^. Due to the small sample volume and the spatial heterogeneity of cutaneous blood flow, LDF measurements of microvasculature ED can suffer from poor reproducibility ^19,20^.

Apart from the abovementioned 1D measurement techniques, imaging has also been employed for ED testing. For example, Laser Doppler imaging (LDI) provides 2D perfusion maps of skin areas (≈ 50 cm^2^) at depths of 1-1.5 mm ^19,20^. However, LDI requires about 1 min to obtain an image, which makes it difficult to track dynamic phenomena during PORH tests ^21^. Laser speckle contrast imaging (LSCI) provides an index of blood flow, yet with a poor penetration depth of ≈ 300 μm, which only allows perfusion measurements of the superficial skin areas ^13,22^. These sensing and imaging techniques can only provide a rough assessment of microvascular function within whole sampled tissue volumes, without directly visualizing the real changes occurring within the microvascular network (e.g., diameter increase, vessel recruitment etc.) during the PORH process [15, 25]. Such detailed visualizations of skin microvascular changes would enable a paradigm shift from arbitrary perfusion/flow recordings from entire tissue/skin areas towards a more quantified ED assessment based on absolute changes of parameters, such as the microvascular diameters and numbers within different skin layers [13].

Recently, Dynamic Optical Coherence Tomography (D-OCT) has been reported to provide detailed visualizations of skin microvasculature and assess endothelial function by measuring absolute changes of microvascular diameter/density, as well as blood flow rate and speed ^23,24^. However, relevant D-OCT studies only assess the microvascular changes with low temporal resolution, without fully resolving and quantifying the fast dynamic changes occurring in the skin microvasculature during the PORH process. In addition, D-OCT provides limited penetration depths (500 μm) in the highly scattering skin tissue, which only allows to assess the superficial dermal vessels ^23^.

Considering the limitations of all abovementioned techniques, there is a need for novel approaches capable of visualizing and monitoring the dynamic changes taking place within the skin microvasculature during functional ED tests with: i) high temporal resolution, ii) high spatial resolution and iii) whole-skin depth analysis capabilities.

Herein, we hypothesized that Fast Plane Raster-Scan Optoacoustic Mesoscopy (FP-RSOM), a further development of RSOM technology which can provide whole-skin depth microvascular imaging at high resolutions ^25,26^, can overcome these limitations by defining new image-based biomarkers to characterize endothelial function and dysfunction. To this end, we introduce a novel approach for the non-invasive assessment of whole-skin microvascular endothelial function by means of FP-RSOM which enhances the unprecedented capabilities of RSOM technology with high temporal resolution (1Hz). We conducted standardized PORH challenges in healthy volunteers and in subjects with high cardiovascular risk due to smoking habits, or with previously diagnosed cardiovascular disease. In particular, we first performed RSOM imaging during PORH challenges in healthy volunteers, allowing comprehensive evaluation of the RSOM capabilities to image dynamic changes in skin microvasculature by measuring absolute quantitative changes of individual vessel diameter and the total blood volume. We next applied FP-RSOM imaging (1 frame/second) to record the fast-scale dynamics of skin microvessels at whole-skin depths during the PORH process and determined several biomarkers, such as the maximum volume change (MVC), hyperemia ratio (HR) and the time to peak (TP), to enable the quantification of the endothelial function of skin microvasculature by quantifying the vessel density changes at various skin depths. Lastly, we benchmarked our FP-RSOM-based method by comparing the microvascular ED between age-matched healthy volunteers and smokers, as well as another group of age-matched healthy volunteers and patients with CVD (n = 10 for each group). We showcase for the first time the markedly different impairments to the endothelial function of upper and deeper dermal microvessels in the smoking group. We validated the FP-RSOM measurements with macrovascular ultrasound-based measurements of the radial artery diameter, as well as microvascular LDF and tissue spectrometry (O2C^©^) measurements of skin blood flow, haemoglobin (Hb) and oxygen saturation (SO_2_). FP-RSOM provided higher sensitivity in detecting the effects of smoking and CVD on microvascular ED compared to LDF and tissue spectrometry.

Altogether, our results demonstrate the great potential of FP-RSOM for elucidating the morphology, functional state and reactivity of skin microvasculature via the definition of novel FP-RSOM-based biomarkers, with great implication for investigating new aspects of endothelial function and its impairment in cardiovascular risk and disease groups.

## Results

### FP-RSOM assessment of skin microvascular endothelial function

In order to assess skin microvascular endothelial function, we employed FP-RSOM and validated our measurements by means of the commercial O2C (Oxygen to See; LEA Medizintechnik, Giessen, Germany) to monitor the reactive hyperaemia process (see Methods) within the skin forearm during a standardized PORH (Fig. 1a, b). The PORH procedure (Fig. 1b) includes three sets of measurements: 2 minutes of baseline, 5 minutes of “cuff on” (inflated cuff at a pressure of at least 40 mmHg above the systolic blood pressure of the subject) and 5 minutes of “cuff off” (cuff deflation). The 3D RSOM scan recorded anatomical and vascular features of the skin (4 × 2 mm^2^) within a minute before the PORH process, then FP-RSOM imaging was implemented by scanning the RSOM head back and forth along a same line (4 mm long) inside the field of view of the 3D scan with a frame rate of 1 Hz. Cross-sectional volumetric RSOM images of the forearm skin for each subject could be used to identify the optimal measurement position by providing detailed visualizations of the microvasculature within the epidermis and dermis layers (Fig. 1c-e). The hyperaemia response of skin microvessels during the total 12 minutes of the arterial occlusion process can be recorded by 3D RSOM or FP-RSOM, with a simultaneous O2C measurement and clinical ultrasound for assessing macrovasculature ED ^27^. Cross-sectional image of the 3D volume acquired at the forearm skin from a healthy volunteer (Fig. 1c) revealed the epidermis and dermis layers at a depth of about 1.5 mm at the illumination wavelength of 532 nm. Maximum intensity projection (MIP) images of the 3D volume depicted the superficial skin ridges and capillary loops (Fig. 1d) and the dermal vasculature in the vascular plexus (Fig. 1e). Inspection of the raw optoacoustic signals of the first FP-RSOM scan provided an estimation of the acquired signal quality (Fig. 1f) in order to proceed with the functional FP-RSOM measurement during the PORH. The reconstructed image of the FP-RSOM scan (Fig. 1g) resolved skin epidermis and dermis layer structures, correlating well with the 3D RSOM image (Fig. 1c). In order to investigate the different responses of microvasculature ED along the skin depth, the dermal vasculature can be separated into the microvessels in the subpapillary dermis (SD) layer and larger vessels in the reticular dermis (RD) layer. We next investigated the obtained 3D RSOM and FP-RSOM data in more detail.

**Fig. 1.**
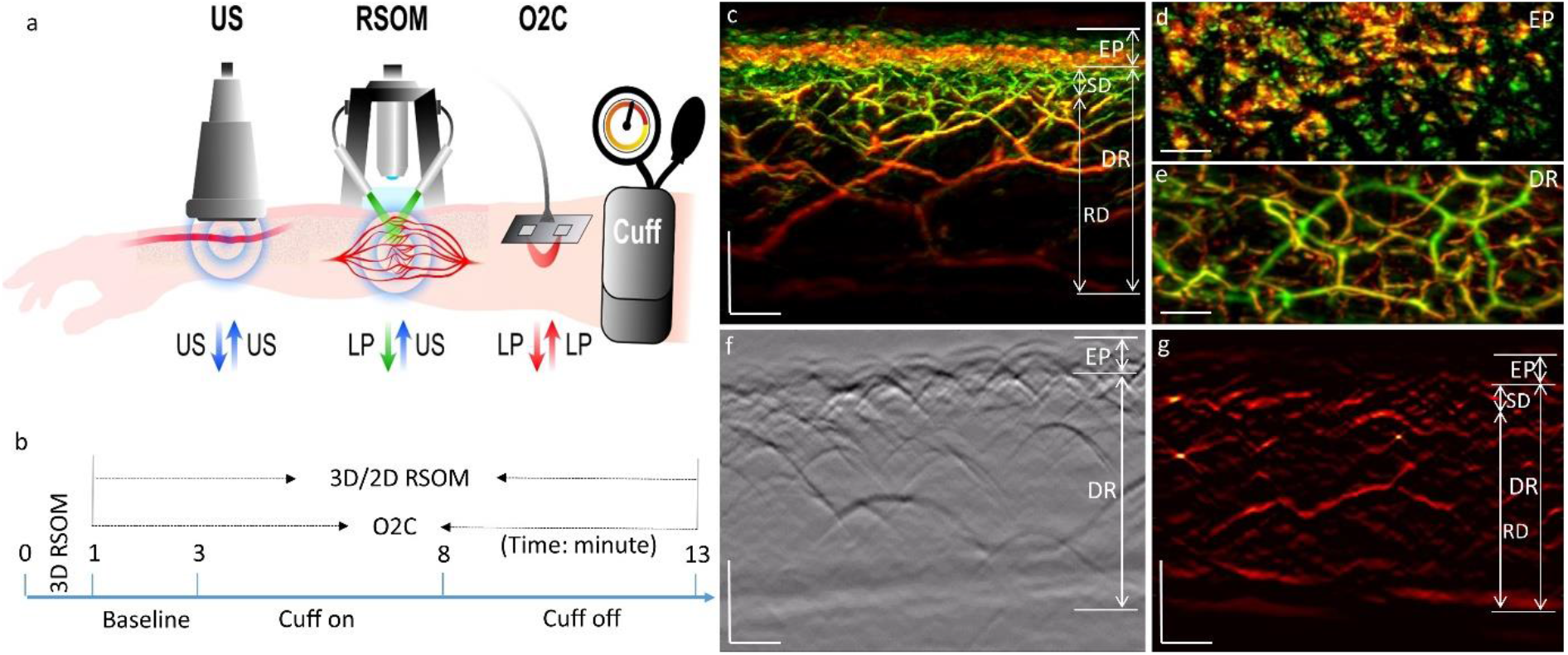
FP-RSOM in assessment of skin microvasculature endothelium function. a, Schematic illustration of assessment of skin microvessel endothelium function at the forearm by FP-RSOM, O2C and ultrasound (US) during PORH test; O2C: oxygen to see, a commercial system including laser Doppler flowmetry (LDF) and White-Light Spectroscopy (WLS) probes, simultaneously recording blood flow, partial blood volume (rHb) and oxygen saturation (SO_2_). b, The timeline of the arterial occlusion assessment by 3D/ FP-RSOM and O2C simultaneously, including the following measurements: 2 minutes baseline, 5 minutes after inflating the cuff on and 5 minutes after deflating the cuff off. c, Cross-sectional image of the 3D RSOM scan at the forearm of a healthy volunteer, where the dermis vasculature (DV) can be grouped into microvessels in the subpapillary dermis (SD) layer and vessels in the reticular dermis (RD) layer. d,e, Corresponding MIP images of the epidermis (EP) and dermis (DR) layers of (c) in the coronal direction. f, Raw optoacoustic signals of one FP-RSOM scan while the corresponding reconstructed image is shown in (g), where the dermal vascular plexus (VP) can be divided into upper plexus (UP) and lower plexus (LP). Scale bar: 500 μm.

### 3D-RSOM visualization of skin microvasculature hyperaemia

In order to assess the performance of RSOM in visualizing the hyperaemia response of skin microvasculature during the PORH, a rectangular skin area (4 ×2 mm^2^) of the forearm of one healthy volunteer was imaged by 3D RSOM at 1-min intervals and the results are presented in Fig. 2. The first 10 cross-sectional RSOM images and corresponding MIP images of the dermis (DR) vasculature in the coronal plane (Fig. 2a-j) clearly identified significant changes of skin microvasculature during the hyperaemia process (*see Supplementary movies 1 and 2*). RSOM images at the baseline (Fig. 2a, b) resolved rich vasculature in the DR layer. After constricting the arteries by inflating the cuff, microvessels in the subpapillary dermis (SD, white arrows in Fig. 2b) layer first started to disappear (Fig. 2c) and became almost completely invisible after 2 minutes (Fig. 2d), while the melanin features in the epidermis (EP) layer remained constant. Afterwards, the vessels in the deeper reticular dermis (RD) layer gradually became invisible (Fig. 2c-g) and the image intensity dropped significantly when comparing the baseline image (Fig. 2b) to the image (Fig. 2g) of the last minute of cuff inflation. A strong hyperaemia response was recorded after deflating the cuff: vascular features were fully recovered and more cutaneous vessels (white arrows in Fig. 3h) were identified. The dilation of three vessels (marked by the white arrow labels 1, 2 and 3 in Fig. 2a) in the hyperaemia process was clearly quantified (Fig. 2m) by measuring the vessel diameter at every minute, exhibiting different response patterns to the pressure stimuli. For example, the mean diameter of vessel 1 was 58 μm at baseline and increased to 69 μm at the peak point of hyperaemia response (18% increment of the vessel diameter) after cuff deflation. The mean diameter of the vessel 2 was 27 μm at baseline and dilated up to 33 μm at the peak point of hyperaemia response (22% increment of the vessel diameter) after cuff deflation. Furthermore, the hyperaemia process (see Methods) was characterized based on the changes of skin total blood volume computed from 3D RSOM images (see Methods) as shown in Fig. 2k, where the total blood volume of the SD, RD and the whole dermal vasculature (DV including SD and RD layer together) were computed separately. The changes of the total blood volume in the SD, RD and DV showed similar patterns and correlated well with the image changes of Fig. 2a-j. The hyperaemia was assessed simultaneously by the O2C® system, measuring changes of blood flow, oxygen saturation (SO_2_) and partial blood volume (Fig. 2l). Therefore, 3D RSOM imaging for the first time enables to monitor and quantify the cutaneous microvasculature changes at signal vessel resolution during the hyperaemia process.

**Fig. 2.**
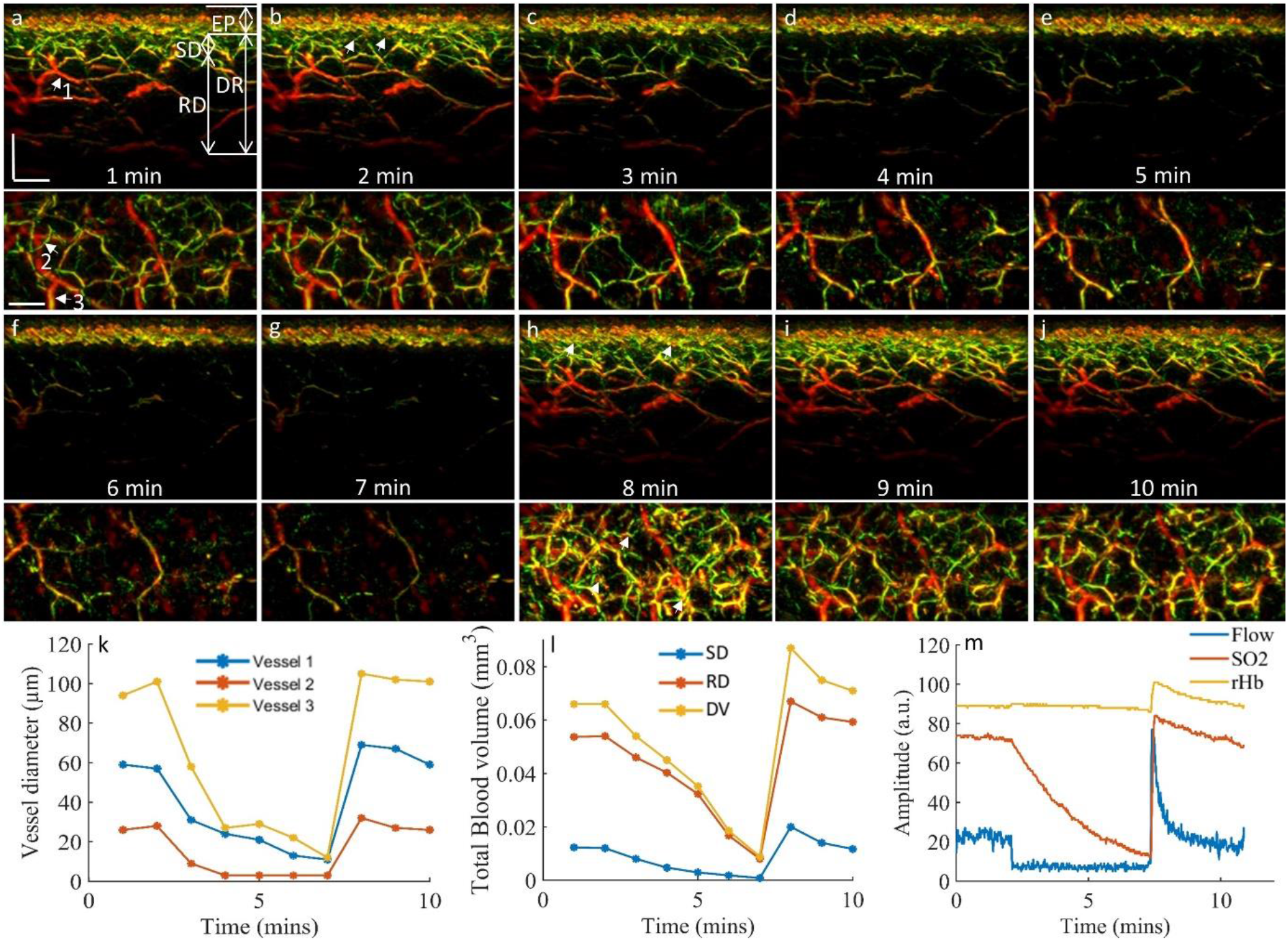
3D RSOM imaging of skin microvasculature hyperaemia during the PORH. The skin vasculature at the forearm (4 ×2 mm^2^) of a healthy volunteer was measured by RSOM at every minute during the 12 minutes PORH process. 10 cross-sectional RSOM images of the first 10 minutes and corresponding MIP images of the dermis layer in the coronal direction (below each cross-section image) are shown in a-j. The white arrows in (h) indicates the new capillaries layer and dermal vessels induced during the reactive hyperaemia process. k, The diameter changes of three vessels (white arrow labels 1, 2 and 3 in a) characterized as the FWHM (full width half maximum) values during the occlusion process. l, Changes of the total blood volume in the subpapillary dermis (SD) layer, in the reticular dermis (RD) layer and the whole dermis vasculature (DV) computed from the first 10 3D RSOM images during the arterial occlusion. m, O2C results including the blood flow, oxygen saturation (SO_2_) and partial blood volume (rHb). Scale bar: 500 μm.

**Fig. 3.**
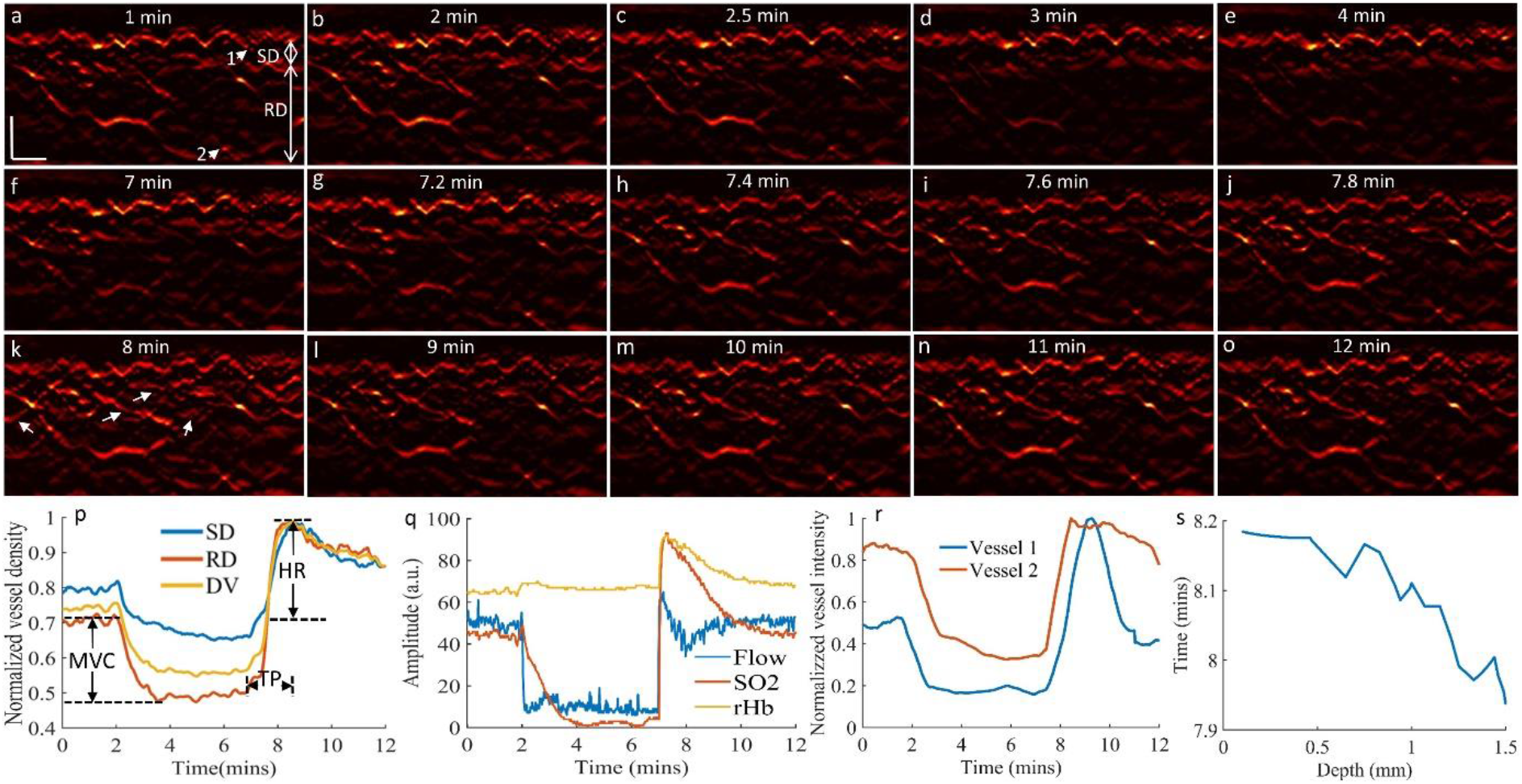
FP-RSOM imaging to quantify skin microvasculature hyperaemia during the PORH. 15 FP-RSOM images at different time points during the PORH are shown in a-o. The white arrows in (k) indicates the new capillaries and dermal vessels induced during the arterial occlusion process. p, Normalized vessel density changes in the subpapillary dermis (SD) layer, in the reticular dermis (RD) layer and the whole dermis vasculature (DV) during the arterial occlusion process. q, O2C results including the blood flow, oxygen saturation (SO_2_) and partial blood volume (rHb). r, Normalized vessel intensity profiles of vessels 1 and 2 [labelled by the white arrows in (a)]. s, Time points of the peak hyperaemia response of skin features at different depths. Scale bar: 500 μm.

### FP-RSOM quantification of skin microvasculature endothelial function

3D RSOM imaging takes about one minute to record rich vasculature image of a skin area of 4 mm × 2 mm, which hinders efficient tracking of the dynamic response of the hyperaemia process. To address this limitation, we implemented FP-RSOM to record RSOM images of a thinner skin volume (4 mm × 0.1 mm), achieving much higher temporal resolutions from one image/minute to one image/second. 15 selected FP-RSOM images of skin features from a healthy volunteer at different time points during the PORH clearly resolved dynamic changes of microvasculature in the subpapillary dermis and reticular dermis layers (*see Supplementary movie 3*). Fig. 3a-e show the skin images at the two-minute baseline time period (Fig. 3a-b) and the first two minutes after inflating the cuff (Fig. 3c-e). In full agreement with the pilot 3D RSOM measurement presented before, the dermal vasculature gradually disappeared while the intensity of melanin features in the epidermis layer remained constant. After deflating the cuff, the hyperaemia response of skin vasculature in the first minute (Fig. 3f-k) was abrupt and prominent, appearing in both the SD to RD layers. New vessels (indicated by white arrows in Fig. 3k) were identified at the peak hyperaemia point. Fig. 3l-o further depicts the recovery response of skin microvasculature after the peak hyperaemia point. The normalized vessel density profiles (see Methods, Fig. 3p) of the SD, RD and whole dermal vasculature (DV) quantified the microvasculature changes during the cuff occlusion process, which correlated well with images in Fig. 3a-o. Besides FP-RSOM imaging, the blood flow, oxygen saturation (SO2) and partial blood volume (rHb) were simultaneously recorded by O2C (Fig. 3n).

In order to quantify endothelial function, we computed three parameters: maximum volume change (MVC), hyperaemia ratio (HR) and the time to peak (TP) from the normalized vessel density profiles of FP-RSOM images and O2C® measurements as illustrated in Fig. 3p (*see Methods*). It should be noted that the MVC and HR values of the RD layer in the hyperaemic state (Fig. 3p) were much higher than of the SD layer, demonstrating differential hyperaemia response of the two different skin vascular beds (SD/RD). Moreover, the vessel density change computed from the FP-RSOM images (Fig. 3p) showed a slower hyperaemic response (about 1 minute from the time of releasing the cuff to the time of the highest peak value) compared to the fast change O2C® measurement (about 10 sec, Fig. 3q). The partial blood volume (rHb) of the deeper vasculature assessed by WLS of O2C® (Fig. 3q) depicted little variations when inflating the cuff while the FP-RSOM profiles relating to the superficial skin microvasculature (Fig. 3p) showed significant decrement as visualized in the FP-RSOM images (Fig. 3a-e). In addition, the normalized vessel intensity profiles of two different vessels (Labels 1 and 2 in Fig. 3a) shows that the microvessel from the SD layer (Label 1 in Fig. 3a) responds faster compared to the deep larger vessels from the RD layer (label 2 in Fig. 3a) when inflating the cuff, while the deep dermal vessel reached the peak of hyperaemia response earlier than the microvessel from the superficial SD layer after deflating cuff. Furthermore, the time of peak hyperaemia response along the skin depth was computed as shown in Fig. 3s, showing an approximately 0.25-minute time difference from the superficial microvessels to the deep dermal vasculature at a maximum depth of 1.5 mm.

### Quantification of smoking effects on skin microvasculature endothelial function

In order to demonstrate the capability of FP-RSOM imaging on studying skin microvasculature endothelial function in subject with increased cardiovascular risk, 10 non-smoking volunteers and 10 age-matched smokers were measured by FP-RSOM during the 12-minute PORH process, while O2C® signals were again simultaneously recorded. Fig. 4a-d shows two cross-sectional MIP RSOM images of the skin at the forearm of a non-smoking male volunteer (Fig. 4a) and a male smoker (Fig. 4b), and two corresponding FP-RSOM images (Fig. 4c and 4d), respectively. The normalized vessel density profiles computed from the FP-RSOM images during the PORH are shown in Fig. 4e (non-smoking volunteer) and Fig. 4f (smoker), while the corresponding O2C® results are depicted in Fig. 4g and 4h, respectively. Our results show good temporal agreement between the FP-RSOM and O2C® readouts. The MVC values (Fig. 4i) were computed (*see Methods*) to measure the maximum blood volume decrease after cuff inflation and vascular occlusion. The MVC values of the smoker group were 0.34 ± 0.08 versus 0.48 ± 0.07 at DV layer, 0.25 ± 0.12 versus 0.44 ± 0.07 in the SD layer and 0.38 ± 0.06 versus 0.48 ± 0.04 in the RD layer compared to the non-smoking group. An unpaired t-test and a Wilcoxon signed-rank test showed significant differences (*P* < 0.001 in DV, P < 0.01 in SD and P < 0.001 in RD, respectively). Furthermore, we found that the MVC value (0.25 ± 0.12) of the SD layer were markedly lower than the value (0.38 ± 0.06) of the RD layer in the smoker group (P < 0.01) while there was no significant difference in the non-smoking group.

**Fig. 4.**
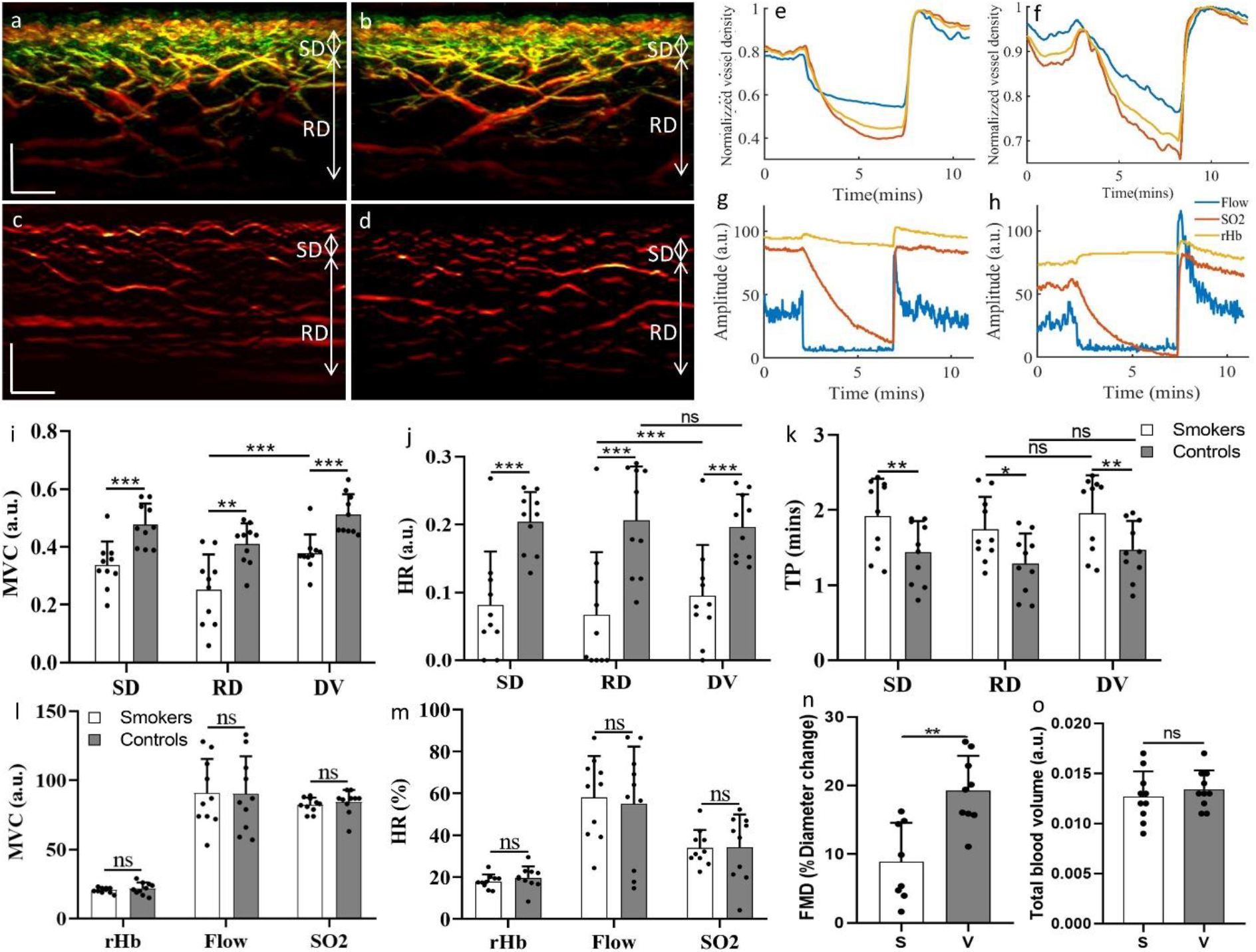
Quantification and comparisons of skin microvasculature endothelial function between non-smoking volunteers and smokers. Cross-sectional RSOM images of a non-smoking male volunteer (a) and a male smoker(b), while corresponding FP-RSOM images are shown in (c) and (d), respectively. e,f, The normalized vessel density profiles used to characterize endothelial function during PORH process between the non-smoking volunteer (e) and the smoker (f). g,h, Corresponding results of the O2C measurements, showing the blood flow, partial blood volume (rHb), and oxygen saturation (SO_2_) values. i-k, Comparisons of the MVC (i), HR (j) and TP(k) values computed from the microvasculature in the subpapillary dermis (SD) layer, the reticular dermis (RD) layer and the whole dermis vasculature (DV) between the non-smoking volunteers and smokers. l,m, Results of MVC and HR parameters calculated from the O2C measurements. n, Ultrasound FMD results characterized by the vessel diameter change between the non-smoking volunteers (V) and smoker (S) groups. o, Comparisons of the total blood volume in the dermal layer and epidermal thickness between the non-smoking volunteers (V) and smoker (S) groups. Scale bar: 500 μm.

The HR values (Fig. 4j) were calculated as a second parameter to quantify the maximum blood volume change ratio (see methods) during the hyperaemia process after deflating the cuff. The smoker group exhibited significant lower values of 8.15 ± 7.89 (%) versus 20.38 ± 4.43 (%) at DV layer, 6.67 ± 9.26 (%) versus 20.63 ± 7.94 (%) in the SD layer, and 5.34 ± 7.50 (%) versus 19.65 ± 4.84 (%) in the RD layer, compared with the non-smoking group. An unpaired t-test and a Wilcoxon signed-rank test showed significant differences (P < 0.001 in DV, P < 0.001 in SD and P < 0.001 in RD, respectively). Moreover, HR values of the SD layer in the smoker group were lower than the values of the RD layers (P < 0.05) while there was no statistical difference in the non-smoking group.

The TP values (Fig. 4k) related to hyperaemia response speed were computed as the time difference from the point of deflating the cuff to the hyperaemia response peak, which also exhibited a marked difference between the smoking and non-smoking groups. The mean TP values of the smoker group were 1.92 ± 0.49 minutes versus 1.44 ± 0.41 minutes of the non-smoking group at DV layer, 1.74 ± 0.43 minutes versus 1.39 ± 0.49 minutes in the SD layer, and 1.96 ± 0.50 minutes versus 1.47 ± 0.38 minutes in the RD layer, respectively. An unpaired t-test and a Wilcoxon signed-rank test showed significant differences (P < 0.01 in DV, P < 0.05 in SD and P < 0.01 in RD, respectively). There was no significant difference of TP values between SD and RD layers in both smoking and non-smoking groups.

As a validation, the MVC (Fig. 4l) and HR values (Fig. 4m) of O2C® measurements were computed, which however exhibited no significant difference between the smoking and non-smoking groups. In addition, the macrovascular endothelial function assessed by clinical ultrasound means during the PORH process also demonstrated significantly changes in the smoking (S) group compared to the non-smoking volunteer groups (V) as shown in Fig. 4n. Furthermore, the total blood volume (Fig. 4o) related to the skin microvasculature morphology features showed no marked differences between the smoking (S) and non-smoking volunteer groups (V).

### Quantification of skin microvasculature endothelial function in patients with CVD

To further showcase the promising capability of the FP-RSOM technology to precisely assess ED, we performed targeted measurements in patients with cardiovascular disease (CVD). 10 CVD patients were measured by FP-RSOM scanning during a 12-minute PORH and compared with 10 healthy volunteers recorded in Fig. 4. Fig. 5a and 5b show two cross-sectional MIP RSOM images of the skin at the forearm of a healthy volunteer and a CVD patient. We found that the MVC values of the CVD group exhibited significant lower values of 0.16 ± 0.048 versus 0.48 ± 0.73 in the healthy volunteer group (P < 0.001). Similarly, the HR values of the CVD group were 7.13 ± 3.41 (%) significantly lower than in the healthy volunteer group 20.38 ± 4.43 (%) with P < 0.001. The TP values (Fig. 5g) of the CVD group were 2.18 ± 0.20 minutes versus 1.44 ± 0.41 minutes in the healthy volunteer group (P < 0.001).

**Fig. 5.**
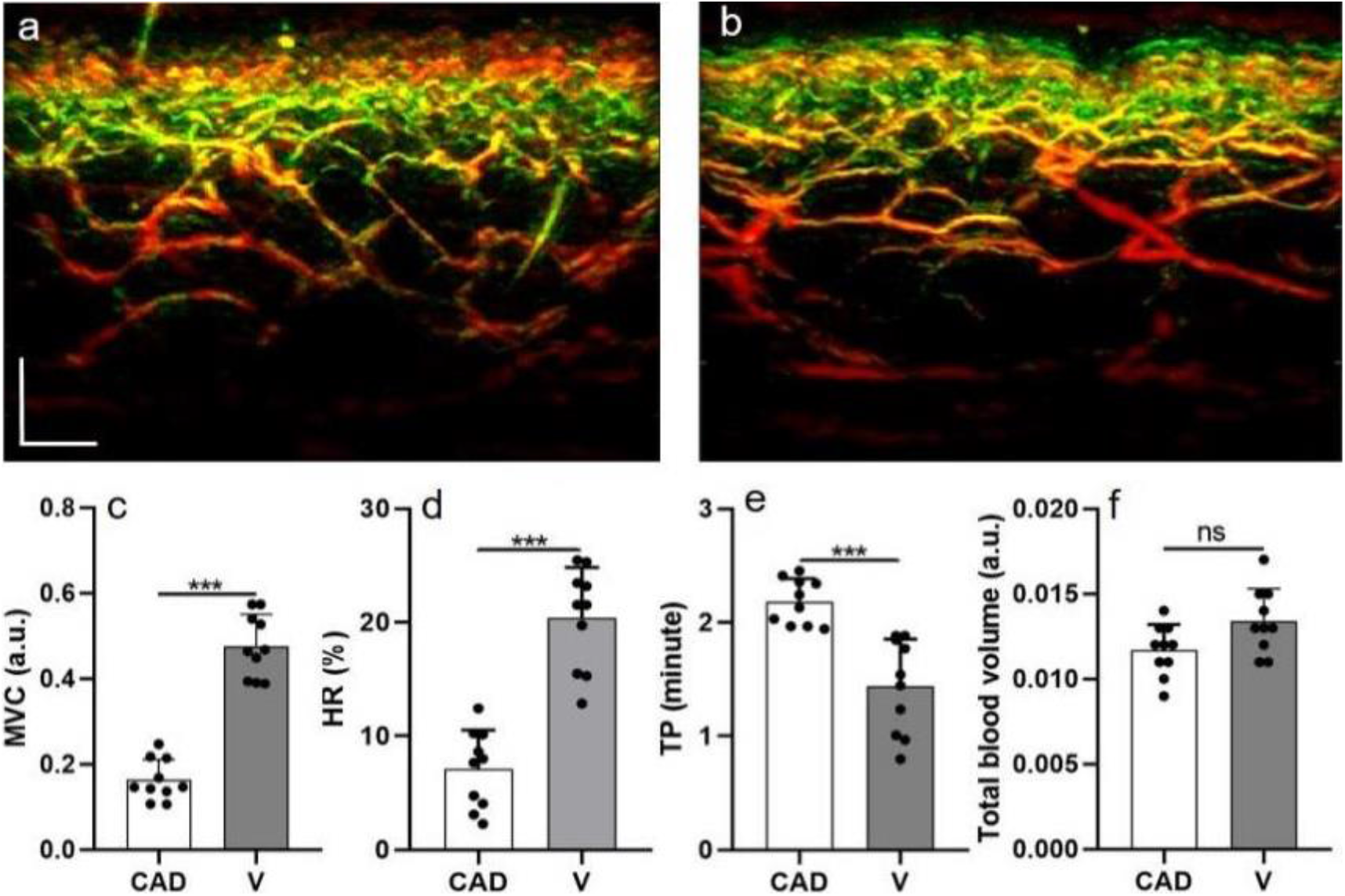
Comparisons of skin microvasculature endothelial function between and healthy volunteers (V) and CVD patients (CVD). a,b, Cross-sectional RSOM images of a healthy volunteer (a) and a CVD patient(b). e-f, Comparisons of the MVC (e), HR (f) and TP(g) values in all skin depth computed from the RSOM image intensity profiles between the healthy volunteers and CVD patients. h, Comparison of the total blood volume in the dermal layer between the healthy volunteers and CVD patients. Scale bar: 500 μm.

## Discussion

Accurate assessment of skin microvascular structure and function is essential in the investigation of the pathophysiology of CVD and stratification of cardiovascular risk. In this work, we employed 3D RSOM imaging for the first time to visualize the responses of cutaneous microvasculature at single-vessel resolution and quantify changes of vessel diameter and skin total blood volume during standardized PORH tests. Furthermore, we developed a novel technique, termed FP-RSOM imaging, that can be used to quantify the endothelial function of the cutaneous microvasculature by computing three biomarkers from the skin vessel density changes profiles derived from FP-RSOM images during functional tests along the whole skin depth. This newly-developed method identified for the first time the differential hyperaemic responses between the microvasculature from the superficial SD layer and deep RD layers. We also benchmarked our novel method in situations with known endothelial dysfunction, in patients with an increased cardiovascular risk (smokers) and in patients with diagnosed CVD, confirming the expected cutaneous microvasculature endothelial dysfunction in both patient groups. Finally, we validated our RSOM measurements by means of US, LDF and WLS, demonstrating superior sensitivity for detection and quantification of skin microvasculature ED in smokers and patients with CVD compared with purely optical methods.

The RSOM approach we implement in this study overcomes many of the limitations associated with currently used techniques. RSOM is the only technique available that can non-invasively provide highly detailed skin features of optical contrast through the entire skin depth. Relying on the 3D RSOM imaging, we for the first time present a comprehensive visualization of skin microvasculature changes in response to pressure stimuli at single-vessel resolution and assess absolute quantitative measures of vessel diameter and the total blood volume through the entire skin depth during the PORH process. The dilation of individual microvessels of varying size located at different skin depths during hyperaemia was precisely measured by 3D RSOM as shown in Fig. 2k as well as the skin total blood volume changes shown in Fig. 2l.

The current implementation of 3D RSOM imaging takes about one minute to scan a field of view of 4 mm × 2 mm, which is too slow to record the fast dynamics of skin microvessels during a PORH. To address this, we implemented FP-RSOM imaging to record the microvasculature responses at 1 Hz of a thin volume (4 mm × 0.1 mm) and quantify the microvasculature endothelial function by computing the skin vessel density change profiles from FP-RSOM images as shown in Fig. 3. By using FP-RSOM, we for the first time quantified the different endothelial function responses of the microvessels located in the superficial subpapillary dermis layer and larger vessels from the reticular dermis layer and found that the time points of the peak hyperaemia response varied along the skin depths. Moreover, comparisons between FP-RSOM, Laser Doppler and WLS as shown in Supplementary figure XXX demonstrated that FP-RSOM can achieve more reproducible results compared to the point measurements of Laser Doppler and WLS.

Smoking is known to be associated with an acute impairment of the microvascular function [34, 35]. We investigated the effects of smoking on skin microvasculature endothelial function by FP-RSOM, validated by Laser Doppler and WLS measurements as well as the microvasculature ED assessed by clinical ultrasound. The MVC and HR values computed from the FP-RSOM vessel density change profiles showed significant differences between the smoker and healthy non-smoker groups while results of the blood flow, partial blood volume (rHb), and oxygen saturation (SO2) values did not detect ED differences. This can be explained by the regional heterogeneity of skin perfusion in the forearm, which leads to spatial variability and contributes to the relatively poor reproducibility of the point sensing techniques (Laser Doppler and WLS); FP-RSOM imaging however records optoacoustic signals over a thin volume (4 mm × 0.1 mm), minimizing the spatial variability. Furthermore, we for the first time quantified that the ED of the microvessels located in the superficial subpapillary dermis layer was significantly more severe compared to the ED of larger vessels from the reticular dermis layer in smoker group as shown in Fig. 4i and 4j. The TP values showed that smoking significantly impaired the vessel recovery speed in the PORH compared to non-smoking healthy volunteers as illustrated in Fig. 4k. The macrovascular ED characterized by the artery diameter changes using clinical ultrasound during the PORH also presented significant differences between the smoking and non-smoking group, correlating well with previous studies [36, 37]. In addition, smoking did not appear to alter the forearm skin microvascular structure in this study, as the total blood volume computed from the 3D RSOM images showed no significant differences, suggesting that the cutaneous microvasculature ED may precede the structure changes. Besides smoking, we further investigated the effects of CVD on CMivED and found that the FP-RSOM biomarkers of CVD patients were significantly lower than in the healthy volunteers, while the structural changes of skin vessels in patient forearms were not obvious.

The endothelial dysfunction in the large conduit arteries and resistance small vessels have been reported to have different correlations with cardiovascular risk factors and may be unrelated to one another, suggesting differential regulation of endothelial function in these two vascular beds ^8,17,18,28,29^. The discrepancy between macro- and microvascular dysfunction could be explained by the fact that they may mirror different stages of disease ^8,30^. The endothelial dysfunction in conduit arteries can reflect a process of existing CAD, while microvascular dysfunction may be a prognostic biomarker, which may precede endothelial impairment in large arteries and the subsequent clinical manifestations ^8-10,30^. For example, several studies have highlighted the detection of microvascular dysfunction as potentially clinically relevant for the early diagnosis of microvascular disorders, before macrovascular disease^11,31,32^. We identified both cutaneous microvasculature ED and macrovasculature ED simultaneously in the smoking group, correlating well with previous findings ^33^. Longitudinal studies on individuals with early stages of CVD diseases are needed to further demonstrate the predictive value of microvascular ED compared to macrovascular ED.

In the current study, the FP-RSOM was implemented by scanning the RSOM probe back-forth along a 4 mm line, recording signals from a thin volume. In order to further enhance the signal quality and minimize the effects of tissue heterogeneity, FP-RSOM can be implemented to scan along multi-lines in a larger field of view of human skin while maintaining the scanning speed by using higher repetition rate of laser source. In addition, the cutaneous microvasculature ED was computed based on the vessel density changes derived from FP-RSOM images related to the haemoglobin concentration changes. Multi-spectra FP-RSOM could be implemented to further investigate the oxygen saturation changes during PORHs, which could provide more information to characterize the microvasculature ED. A thorough comparison between FP-RSOM and O2C® is challenging, taking into account the different field-of-views of the two technologies. Furthermore, it is not possible to clearly define the tissues measured by O2C since no tomographic images of the examined region are provided. However, FP-RSOM can quantify the cutaneous microvasculature ED at different skin depths while Laser Doppler and NIRS can only provide an overall estimation of skin perfusion changes.

The development and validation of a novel RSOM-based method to quantify skin microvasculature endothelial function presented herein could serve as a valuable early marker for the diagnosis and monitoring of cardiovascular diseases in the near future.

## Methods

### Study design

All participants signed an informed consent before included in the study. Healthy volunteers freely agreed to participate in full accordance with the work safety regulations of Helmholtz Zentrum München. The study was approved by the local ethics committee of the Technical University of Munich. In the current study we included n_1_ = 10 non-smoking volunteers (6 males, 4 females, mean age 32 ± 4 years), n_2_ = 10 smoker volunteers (7 males, 3 females, mean age 34 ± 5 years, mean cigarettes per day 8.1 ± 4.1, smoking history of 13.3 ± 4.1 years) and n_3_ = 10 patients with cardiovascular disease (9 males, 1 female, mean age 65.4 ± 9.9 years). Volunteers were free of other cardiovascular risk factors and not taking any regular medication.

Tests were performed in a quiet room at normal temperature (≈ 24°C). Subjects were asked to lie in supine position with the ventral side of the dominant distal forearm easily accessible. A blood pressure measurement (pneumatic) cuff of appropriate size was placed around the upper arm and stayed there for the whole duration of the PORH. Before the test, subject ‘s blood pressure was measured to check that all subjects were normotensive. Then, an optically and acoustically transparent plastic membrane was affixed over the forearm using tape. The RSOM scanning head, which carried both the illumination output and an ultrasound transducer (UT), was brought close to the membrane in order to position the focal point of the UT slightly above the skin surface and thereby maximize detection sensitivity. The RSOM scanning head was positioned approximately 5 cm proximal to the wrist while the O2C probe was fixed by means of a transparent foil nearby it, both at the ventral side of the forearm. Moreover, for further validating our RSOM measurements, a traditional ultrasound probe for arterial diameter measurements was placed over the distal radial artery (cross-sectional view) for the whole duration of the cuff measurement.

Each cuff measurement lasted 12 minutes, including: 2 min baseline measurement (deflated cuff), 5 min arterial occlusion measurement (inflated cuff at a pressure of at least 40 mmHg more than the systolic blood pressure of the subject) and 5 min rest measurement (deflated cuff). The cuff was manually controlled by an experienced operator and RSOM, O2C and ultrasound recordings were full synchronised throughout the whole test.

### RSOM imaging system and O2C validation

The present study used an in-house built portable RSOM imaging system featuring a transducer with central frequency of 50 MHz (Fig. 1a), which has been described in detail from our previous studies ^25,34^. An optically and acoustically transparent plastic membrane was affixed using surgical tape on the patient ‘s skin. The scanning head containing the laser output and an ultrasound transducer (UT) was brought close to the membrane in order to position the focal point of the ultrasound detector slightly above the skin surface and thereby maximize detection sensitivity as shown in Fig. 1. An Onda laser (Bright Solutions, Italy) with dimensions of 19×10×9 cm^3^ was used to provide light with wavelength of 532 nm. The repetition rate of the laser was up to 1 kHz, yielding an optical fluence of 3.75 μJ/mm^2^ under the safety limit. For 3D RSOM imaging, a field of view (4 mm × 2mm) was scanned with step size 12 μm in the fast axis and 15 μm in the slow axis. The total scanning time of one measurement took about 60 s. As for the FP-RSOM imaging, the RSOM head was moved back-forth along the middle line (4 mm long) in the defined field of view of the 3D scan (4 mm × 2mm) with 500 Hz repletion rate, thus achieving 1 frames/second. For each measurement, the RSOM scanning head was positioned at the area about 5 cm to the wrist while the O2C probe was fixed nearby as illustrated in Fig. 1. A clinical used pneumatic cuff was placed at the level of the upper arm (i.e. distal to the site of brachial artery measurement) and controlled by an experienced operator. The O2C machine is a commercial lightguide tissue spectrophotometry system ^35^ including Laser Doppler flowmetry and White-Light Spectroscopy probes is used to simultaneously record the blood flow, partial blood volume and oxygen saturation during the occlusion-induced hyperaemia for comparisons. The hyperaemia measurement takes in total of 12 minutes, including: 2 minutes ‘ baseline, 5 minutes ‘ cuff on (inflate the cuff pressure to 220mmHg) and 5 minutes ‘ cuff off (cuff deflation). During the cuff measurements, the FP-RSOM scanning was synchronized with the O2C measurements. In order to visualize the skin microvessels during the 12 minutes ‘ cuff measurement, 12 RSOM 3D scans were recorded in an area of 4 mm × 2mm as shown in Fig. 2. FP-RSOM scans were implemented to record the dynamic response of skin microvessels in a line (4 mm length) of skin.

### Image reconstruction for 3D/FP-RSOM imaging

Our motion correction algorithm was first applied to eliminate motion artifacts in RSOM data ^36^. For 3D RSOM image reconstruction, the acquired RSOM signals were resolved into two frequency bands 10-40 MHz (low) and 40-120 MHz (high) for the 10-120 MHz bandwidth. Signals of the two frequency bands were independently reconstructed and combined to produce a final image as described in previous work ^25^. The reconstruction time for each band took about 5 minutes with the voxel size of the reconstruction grid being 12 μm × 12 μm × 3 μm. Finally, analysed RSOM images were subsequently rendered by taking the maximum intensity projections of the reconstructed images along the slow axis or the depth direction as shown in Fig. 1. For FP-RSOM image reconstruction, the acquired RSOM signals were filtered at a bandwidth of 10-120 MHz. Since the ultrasound detector used in RSOM has a diameter of 3 mm and detection aperture of 60 degrees, optoacoustic signals of the FP-RSOM scan are actually an integration of thin volume (4 mm × 0.1 mm). Thus, 3D reconstruction based on a beam-forming algorithm were used to generate a three-dimensional image stack (4 mm × 0.1 mm) and the final reconstructed image was rendered by taking the maximum intensity projections along the slow axis as shown in Fig. 1.

### Biomarker computation to quantify endothelial function and general statistics

The endothelial function of skin microvasculature was quantified based on the RSOM image intensity changes during the occlusion measurements, while the signals of the blood flow, partial blood volume (rHb) and oxygen saturation (SO_2_) were directly read out from the commercial system. The surface of 3D/FP-RSOM images were first flattened. Then the epidermal and dermis layers were separated into two parts based on an in-house automatic segmentation algorithm. The mean values of the RSOM image intensities from the epidermal layer, dermis layer and the full skin depths were computed separately to characterize the different responses to cuff pressure as shown in Fig. 2k and Fig. 3p. As illustrated in Fig. 3p, several parameters were computed based on the profiles of FP-RSOM image intensity changes: mean values of baseline (*MVB*, 0-2 minutes), the mean values of signals when inflating cuff (*MVIC*, 2-7 minutes), the peak intensity value (*PIV*) and corresponding time point after deflating cuff. Three biomarkers were computed to quantify endothelial function, including the maximum volume change, the hyperaemia ratio and the time to peak intensity. The maximum volume change (*MVC*) was computed as the difference between the *MVB* and *MVIC* values (*MVC = MVB-MVIC*). The hyperaemia ratio (*HR*) was calculated as the ratio: (*PIV-MVB*)/*PIV* (%). The time to peak intensity (*TP*) relating to the hyperaemia response speed was calculated as the time difference from the cuff releasing point (end of 7 minute) to the time of the peak intensity value. Besides RSOM, the three biomarkers were computed for the signal profiles of the blood flow, partial blood volume (rHb) and oxygen saturation (SO_2_) as well.

All metrics were displayed into column table with mean value and standard deviations as error bar. To assess the significance of the statistical differences for the metrics between healthy and patient groups, and sub-groups among patients with diabetes, we performed parametric tests (unpaired t test) for normally distributed data; otherwise, nonparametric tests (Mann Whitney U test) were applied. Statistical significance was assumed at P < 0.05.

### Repeatability and reproducibility test

We performed studies to evaluate the repeatability and reproducibility of FP-RSOM for endothelial function characterization while comparing with O2C. The skin endothelial function of three healthy non-smoking young volunteer (2 females and 1 male, average age 29 ± 2 years old) were measured repeatedly by FP-RSOM imaging and O2C during the 10 minutes ‘ arterial occlusion on consecutive three days. Similar areas of the forearm of the three volunteers were measured every day by a common operator with same environment conditions. The image intensity profiles of FP-RSOM and signal profiles of O2C are shown in the supplementary Fig. S5. We found that the variations of FP-RSOM measurements are much less than the profiles of O2C measurements. The three biomarkers computed from each measurement demonstrated the high repeatability of FP-RSOM compared to the O2C results. In addition, the skin endothelial function of nine healthy non-smoking young volunteers (30 ± 3 years) were measured by FP-RSOM imaging and O2C simultaneously during the 10 minutes ‘ arterial occlusion. The image intensity profiles of FP-RSOM and signal profiles of O2C (including Flow, SO2 and rHb) were normalized by subtracting the minimum values. As shown in supplementary Fig. S6, we observed that the variations of the FP-RSOM measurements among healthy volunteers are much less than the O2C measurements. The three biomarkers computed from each measurement were used to characterize the good reproducibility of FP-RSOM compared to the O2C setup.

## Acknowledgements

This project has received funding from the European Union ‘s Horizon 2020 research and innovation programme under grant agreement No 871763 (WINTHER) and No 687866 (INNODERM), from the European Research Council (ERC) under the European Union ‘s Horizon 2020 research and innovation programme under grant agreement No 694968 (PREMSOT) and from Helmholtz Zentrum München through Physician Scientists for Groundbreaking Projects, in part by the Helmholtz Association of German Research Center, through the Initiative and Networking Fund, i3 (ExNet-0022-Phase2-3). We thank Dr. Sergey Sulima for his attentive reading and improvements of the manuscript and the staff at the Clinic of Vascular and Endovascular Surgery, Klinikum rechts der Isar at TUM for assisting with the patient studies.

## Notes

### Competing Interest Statement

The authors have declared no competing interest.

